# Activity-driven proprioceptive synaptic refinement in the developing spinal cord by complement signaling mechanisms

**DOI:** 10.1101/2025.08.22.671861

**Authors:** Chetan Nagaraja, Serena Ortiz, Akash R. Murali, Aiza Sarwar, Theanne N. Griffith

**Affiliations:** Department of Physiology and Membrane Biology, University of California, Davis, Davis, CA, USA; Undergraduate Program in Neurobiology, Physiology and Behavior, University of California, Davis, Davis, CA, USA; Howard Hughes Medical Institute

## Abstract

Proprioceptive group Ia afferents detect muscle stretch to guide effortless and purposeful movement and make monosynaptic connections with spinal a-motor neurons to mediate reflexes, such as the stretch reflex. It is thought that proprioceptive Ia afferents target motor neurons of the same spinal segment; yet, how this specificity, if any, is established during early development is unknown. Using *ex vivo* spinal cord electrophysiology preparations from neonatal mice of both sexes, we identified a developmental period during which proprioceptive la afferents evoke both segmental and intersegmental responses at monosynaptic latencies. We provide anatomical evidence that motor neurons in the lumbar segment 4 (L4) receive direct input from proprioceptive Ia afferents in L5 during early postnatal development. Intersegmental responses (L4/L5) were prominent at postnatal days (P) 4–7 but were virtually absent by P11–13. To test the role of proprioceptor activity on segmental specification, we analyzed Na_V_1.6 conditional knockout mice (Na_V_1.6^cKO^), in which proprioceptor signaling is impaired, and found that intersegmental responses persist up to P11–13 but were absent in age-matched floxed controls. We predict this is due to impaired activation of complement signaling pathways, as Na_V_1.6^cKO^ mice showed reduced C1qA expression in the ventral spinal cord at P9. Consistent with this, C1qA knockout mice also retain intersegmental responses at P11–13. Collectively, these findings identify an important postnatal window during which segmental specificity of proprioceptive circuits emerges and suggest that proprioceptor activity induces C1qA-mediated elimination of excessive intersegmental connectivity.

**Key points summary:**

- During the first ten days of postnatal development, the spinal monosynaptic reflex arc possesses intersegmental proprioceptive afferent projections onto motor neurons of an adjacent segment.
- These exuberant Ia-motoneuron projections are accompanied by evocable intersegmental reflex responses. In normal conditions, the intersegmental response is lost by postnatal day 11. Conversely, in a mouse model of impaired proprioceptive signaling, the intersegmental response persists.
- We find that impaired proprioceptive signaling through the monosynaptic reflex arc leads to low expression levels of the complement signaling protein, C1qA, near ChAT-positive motor neurons, whereas microglia recruitment is unaffected. This was consistent with the persistence of robust intersegmental responses in C1qA knockout mice.
- Collectively, our results show that proprioceptor activity induces complement cascade signaling to prune exuberant intersegmental synapses in the spinal cord formed during early development, resulting in segmental restriction of the monosynaptic reflex arc by the end of the second postnatal week.

## Introduction

The spinal monosynaptic reflex arc comprises stretch-elicited signals from muscles that are conducted along proprioceptive Ia afferents into the spinal gray matter, where they activate motor pools that lead to muscle contraction. As such, this reflexive mechanism prevents excessive muscle stretch and is essential for motor reflexes. The neuroanatomical basis for this reflex arc comprises monosynaptic connections between proprioceptive Ia afferents and α-motor neurons, has been extensively studied, and is a feature that is conserved across species (Eccles *et al*., 1957; Fritz *et al*., 1989; Wenner & Frank, 1995; Hongo *et al*., 1997; Mears & Frank, 1997; Mendelsohn *et al*., 2015; Espino *et al*., 2025). In addition to α-motoneurons, Ia afferent collaterals also make monosynaptic contacts with several other spinal neuron populations, including Ia inhibitory interneurons, intermediate zone interneurons, and dorsal and ventral spinocerebellar tract neurons (Jankowska & Lindström, 1972; Harrison & Jankowska, 1985; Bannatyne *et al*., 2009; Jankowska, 2015). The present study focuses specifically on the Ia-α-motoneuron connection, given its well-defined role in the stretch reflex.

Proprioceptive Ia afferent connectivity is preferential to α-motor neurons innervating the same muscle (homonymous connectivity), while connections to motor pools that innervate muscles performing synergist functions (heteronymous connectivity) are much less prevalent, and connections targeting antagonist muscles are avoided (Eccles *et al*., 1957; Mears & Frank, 1997; Mendelsohn *et al*., 2015). The spinal cord is anatomically divided into segments, with each spinal segment named after its respective doral and ventral roots. While motor pools can span spinal segments, it is known that individual motor neurons and their corresponding ventral roots show segmental specificity (Blivis *et al*., 2019). It is unknown, however, whether the spinal monosynaptic reflex arc possesses similar segmental organization. Are proprioceptive Ia afferents restricted to α-motor neurons of the same spinal segment, or do they target α-motor neurons in neighboring segments? Such insight is important for advancing our understanding of mechanisms underlying spinal cord circuit development.

The developing nervous system is characterized by a period of supernumerary synapses, a phenomenon observed across different nervous system modalities and species (Colman *et al*., 1997; Watts *et al*., 2003; Stevens *et al*., 2007; Schafer *et al*., 2012; Hashimoto & Kano, 2013). Supernumerary synapses are then pruned during a period of synaptic refinement by activity dependent mechanisms mediated primarily by microglia, though some studies report a role for astrocytes as well (Shatz & Stryker, 1988; Stevens *et al*., 2007; Schafer *et al*., 2012; Chung *et al*., 2013). It is unknown whether during development, the monosynaptic reflex arc is also characterized by a period of supernumerary synapses that span spinal segments, followed by a period of synaptic refinement. Both microglia, central nervous system resident immune cells, and the classical complement signaling pathway, an integral part of the innate immune response, have been implicated in physiological synaptic refinement, as well as excessive synaptic pruning during disease (Stevens *et al*., 2007; Schafer *et al*., 2012; Vukojicic *et al*., 2019). Thus, we asked whether similar mechanisms were involved in synaptic refinement of the monosynaptic reflex arc during postnatal development.

To address this, we used a hemisected *ex vivo* spinal cord preparation in which we recorded segmental and intersegmental proprioceptor Ia-α-motor neuron responses across development. By stimulating the fifth lumbar dorsal root (DRL5) and recording from either the fifth or fourth lumbar ventral root (VRL5, VRL4) in C57Bl/6J mice, we observed significant intersegmental responses (DRL5 to VRL4) with monosynaptic latencies at postnatal day 4 (P4). These intersegmental responses persisted until P10 but were nearly absent by P11. Using ventral and dorsal root tracing, we confirmed direct targeting of motor neurons in L4 by proprioceptive afferents from L5, supporting the notion that these intersegmental responses are monosynaptic. Moreover, consistent with prior work, we showed that segmental refinement was dependent upon proprioceptor activity, as we observed a persistence of an intersegmental response in a conditional knockout mouse in which the voltage gated sodium channel, Na_V_1.6, is genetically deleted in all somatosensory neurons (Pirt^Cre^;Na_V_1.6, Na_V_1.6^cKO^, (Espino *et al*., 2025)).

We explored a possible role for microglia and complement signaling in this process. We found no differences in the number of microglia in motor regions of the spinal cord at P9 in Na_V_1.6^cKO^ mice compared to floxed controls. Conversely, at P9 in Na_V_1.6^cKO^ mice compared to controls, we observed a significant decrease in C1qA expression, an important component of the classical complement signaling pathway (Stevens *et al*., 2007). This suggests a role for activity-dependent activation of the complement system for synaptic pruning and segmental specification of proprioceptive axons comprising the monosynaptic reflex arc. Indeed, like Na_V_1.6^cKO^ mice, we found that C1qA knockout mice also retained an intersegmental response at P11. Taken together, our findings demonstrate that segmental specificity of proprioceptive Ia-α-motor neuron synapses require normal electrical activity in proprioceptive Ia afferents, which results in activation of classical complement signaling and pruning of intersegmental synapses.

## Materials and Methods

### Animals

C57Bl/6J mice (strain #000664) and C1qa knockout mice (strain #031675) were purchased from the Jackson Laboratory. Pirt^Cre^ and Scn8a^fl/fl^ mice were gifted by X. Dong (John Hopkins University, (Kim *et al*., 2008)) and M. Meisler (University of Michigan, (Levin & Meisler, 2004)), respectively. All mice used were on a C57BL/6J background (non-congenic). All experiments were carried out in compliance with the University of California, Davis Animal Care and Use Committee (Animal Protocol Number 23049) in accordance with guidelines from the National Institutes of Health’s Guide for the Care and Use of Laboratory Animals. Genotyping was outsourced to Transnetyx Inc. Mice were provided food and water *ad libitum* and maintained on a 12-hour light/dark cycle. Both male and female mice were used in all experiments.

### Ventral root electrophysiology

Postnatal mice ranging from P4 to P18 were used for *ex vivo* ventral root electrophysiology. To harvest the spinal cord, mice were deeply anesthetized with isoflurane, decapitated, and eviscerated. The protocol we follow has been used to record motor activity from mice of weight-bearing age (Jiang *et al*., 1999; Nagaraja, 2020; Espino *et al*., 2025). In brief, the preparation was pinned to a dissecting chamber post-evisceration and continuously perfused with ice-cold solution, comprising: 188 mM sucrose, 25 mM NaCl, 1.9 mM KCl, 10 mM MgSO, 0.5 mM NaH_2_PO_4_, 26 mM NaHCO_3_, 1.2 mM NaH_2_PO4, 25 mM glucose, bubbled with 95% O_2_/5% CO_2_. A ventral laminectomy was performed to expose the spinal cord. The dorsal and ventral roots were isolated over the L4 and L5 segments, bilaterally, just proximal to the DRG. The entire cord with the attached roots was removed from the vertebral column after transecting at the thoracic level (T5 - T8). The spinal cord preparation was then transferred to the recording chamber and continuously perfused with artificial cerebrospinal fluid (aCSF): 128 mM NaCl, 4 mM KCl, 1.5 mM CaCl_2_, 1 mM MgSO_4_, 0.5 mM NaH_2_PO_4_, 21 mM NaHCO_3_, 30 mM d-glucose, bubbled with 95% O_2_/5% CO_2_. A midsagittal hemisection was performed, and the spinal hemicords were allowed to equilibrate in aCSF maintained at ambient temperature. After equilibration, the ipsilateral dorsal root (DRL5), the ipsilateral segmental ventral root (VRL5), as well as the ipsilateral VRL4 root were placed into suction electrodes, as shown in the schematic in Figure 1A. DRL5 was stimulated with a single-pulse stimulus, delivered every 30 s, over 10 trials. The stimulus amplitude was set at twice the threshold intensity of stimulation (2T) and a pulse width of 0.1-ms was delivered using a stimulus isolator unit (A365, World Precision Instruments). Extracellular monosynaptic reflex recordings were made at the segmental (VRL5) and intersegmental (VRL4) ventral roots. The data was acquired using Clampex software (v11.2, Molecular Devices) and saved on a computer for offline analysis. Monosynaptic reflex parameters were extracted from the signals for each experiment after averaging the 10 trials using Clampfit (v11.2, Molecular Devices). Differences in the latencies of onset for pairs of segmental and intersegmental responses were determined, and across experiments the difference was found to be 110 ms ± 130 ms. Signals were filtered between 0.1 and 5000 Hz, amplified 1000 times (Model 1700, A-M Systems), and digitized at 10 kHz using Digidata 1440A.

**Figure 1.**
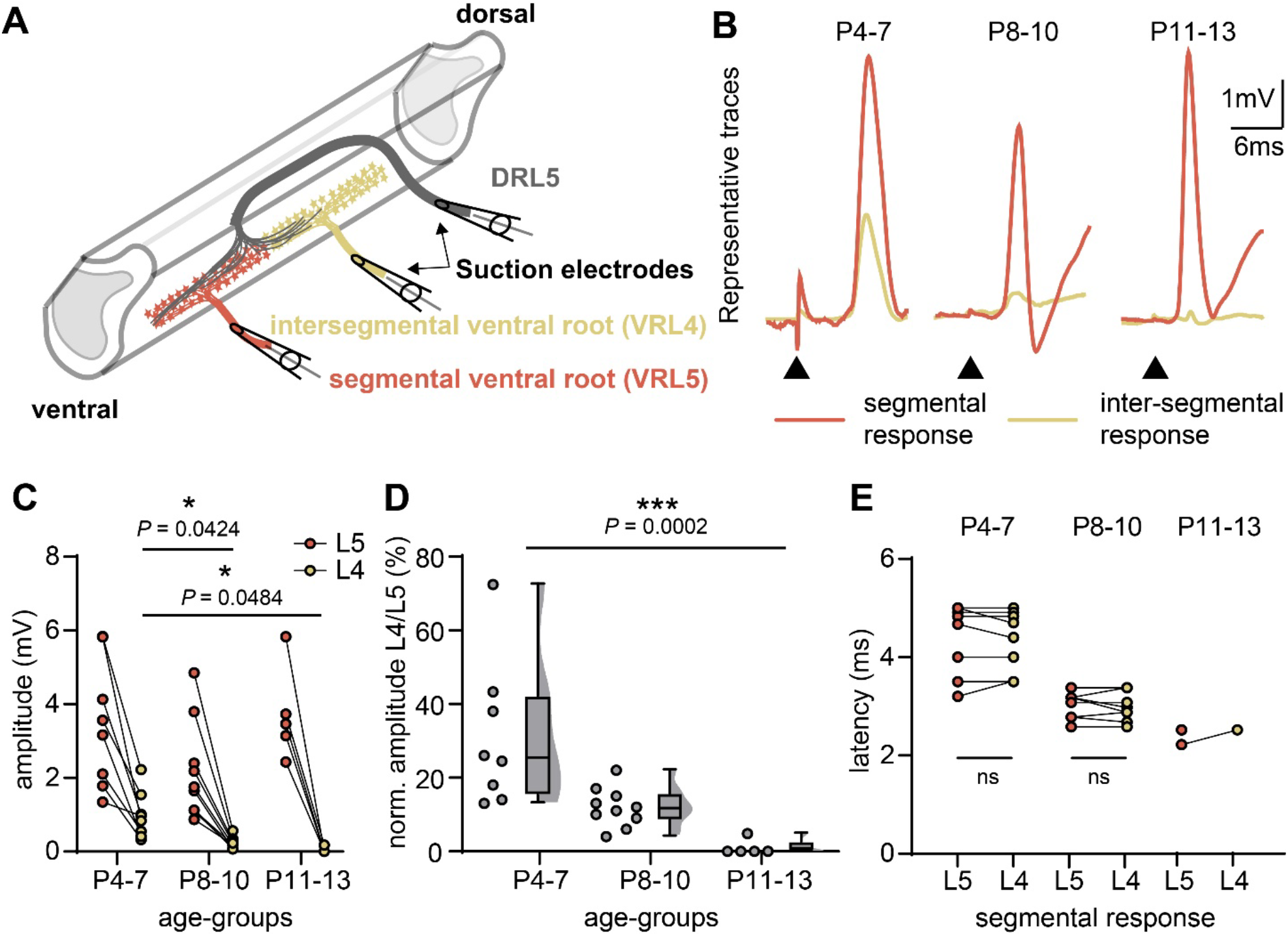
The spinal monosynaptic reflex becomes segmentally restricted by the end of the second week of postnatal development in C57Bl/6J mice. A. Schematic of the *ex vivo* spinal hemicord preparation used for ventral root recordings. Only the L4 and L5 segments are shown for simplicity. DR - dorsal root, VR - ventral root. B. Representative traces of segmental (L5, orange) and intersegmental (L4, yellow) reflex recordings. Each pair represents responses from the same preparation. Arrowheads indicate onset of stimulation. C. Plot showing the peak amplitude of the monosynaptic reflex response. Straight lines connect pairs of segmental (orange dots) and intersegmental (yellow dots) values from the same recording. D. Plot showing the peak amplitude of intersegmental response normalized to the segmental response from each recording. Statistical tests included a two-way ANOVA followed by Tukey’s multiple comparisons test (C) and a one-way ANOVA (Kruskal-Wallis test) followed by Dunn’s multiple comparisons test (D). E. Plot showing the latencies of onset for L5 and L4 response. Straight lines connect pairs of segmental (orange dots) and intersegmental (yellow dots) values from the same recording. Statistical tests included Welch’s t-test for comparison between latencies of onset for L5 vs L4 response: P4-7: *P* = 0.9213, P8-10: *P* = 0.8114, P11-13: intersegmental n too small. C-E. P4-7: n = 8 hemicords, N = 5 animals; P8-10: n=10 hemicords, N = 5 animals; P11-13: n= 5 hemicords, N = 3 animals.

### Tracing of sensory afferents and motor neurons

We visualized proprioceptive afferents and motor neurons by applying fluorescent dextran tracers to cut dorsal and ventral roots as described in Vrieseling and Arber, 2006. We applied a red tracer (Texas Red, 3000MW; Cat. No.: D3328, Invitrogen, converted to yellow for visualization purposes) to the cut end of DRL5 to label L5 proprioceptive afferents and visualize their central projections onto motor neurons. We retrogradely labeled motor neurons of the L4 segment by applying a green tracer (Fluorescein, 3000MW; Cat. No.: D3306, Invitrogen, converted to magenta for visualization purposes) to the cut VRL4 root. The dye was prepared by dissolving the tracers to a final concentration of 1-2% in aCSF. The cords were harvested and hemisected as described earlier and transected at the L3 and L6 segments. The L3-L6 spanning segments of the hemicord preparations were then transferred to a tracing chamber containing aCSF continuously bubbled with 95% O_2_/5% CO_2_. The dorsal and/or ventral roots were placed in tightly fitting suction electrodes and the dye applied to the roots. The cords were left in the oxygenated aCSF overnight for labeling. This was followed by drop fixing in 4% paraformaldehyde (PFA) for 1 hour and storing in 30% sucrose solution overnight. Cryosectioning of the cords was performed as described below.

### Immunohistochemistry

Spinal cord dissections were carried out as described above and lumbar spinal cord spanning L4-L6 segments were harvested and fixed in 4% PFA for one hour. This was followed by washing in PBS (3 washes x 10 minutes per wash), and incubation in 30% sucrose solution overnight at 4°C for cryoprotection. Spinal cords were then embedded in optimal cutting temperature (OCT, ThermoFisher Scientific, no. 4585) and stored in −80°C until cryosectioning. OCT blocks containing the spinal cord were transferred from −80°C to the cryostat and allowed to reach the chamber temperature for an hour. The blocks were then mounted on mounting disks and sectioned at a thickness of 50 mm.

Spinal sections were obtained from male and female P5, P9, and P12 mice. The following primary antibodies were used at the dilutions indicated: rabbit polyclonal anti-Iba1 (1:750; FUJIFILM, 019-19741), mouse monoclonal anti-ChAT (1:200, Millipore Signa, MAB305) and rabbit monoclonal anti-C1q (1:1000; Abcam, ab182451). The following secondary antibodies were used at the dilutions indicated: Goat anti-mouse IgG1 AF647 (1:500, Invitrogen, A21240), Goat anti-Rabbit IgG AF 594 (1:500; Invitrogen, A32740) and Goat anti-Rabbit IgG AF 647 (1:500. Invitrogen, A21244). A hydrophobic barrier was applied to the slides. Sections were incubated for 1 hour in blocking solution (0.1% PBS-Triton-X and 5% Normal Goat Serum), followed by overnight incubation in the primary antibody. After primary antibody incubation, slides were washed 3 times for 5 minutes each in PBS, followed by a 45-minute secondary antibody incubation. Following the addition of a secondary antibody, slides were washed in a dark humid chamber in PBS (5 washes at 10 minutes each). Specimens were mounted with Fluromount-G with DAPI (SouthernBiotech, 0100-20).

### Analysis of microglia numbers and C1qa expression

Spinal sections were imaged using a FV3000 Olympus confocal microscope and either 20X air (0.75 NA), 40X water immersion (0.95 NA), 60X oil immersion objectives (1.4 NA) at 1 μm z-stacks. Images were taken in the ventral horn of the spinal cord where ChAT-expressing motor neurons were located and analyzed in Fiji (ImageJ). Criteria for inclusion of counted microglia included having a soma and one or more emanating processes. For C1qa analysis, each image was thresholded equally to exclude background fluorescence. The mean gray value was calculated to measure the average optical intensity of C1qa fluorescence per image.

### Analysis of proprioceptive inputs on motor neurons

Spinal sections were imaged using a FV3000 Olympus confocal microscope and 60x oil immersion objective (1.4 NA). Z-stacks were obtained with z-step increments of 0.5 mm. We quantified the percentage of L4 motor neurons (magenta) that received 1 or more putative proprioceptive inputs (yellow puncta) on their soma and/or the first 25 mm of proximal dendrites. We included both sexes and carried out the analysis at two age-groups: P4-7 and P8-10.

### Data analysis

Summary data are presented as means ± SD, from *n* hemicords or sections and *N* animals. All analyses of immunofluorescent images contained at least three biological replicates per age and/or genotype. The investigator was blinded to genotypes, sex and age during experimentation and analysis. For all electrophysiological experiments, the investigator was blind to genotype. Prism 10.1 (Graph-Pad software) was used to carry out all statistical tests. Analysis of statistical differences of normally distributed data were determined using parametric tests, and data that did not conform to Gaussian distributions or had different variances were determined using nonparametric tests. Statistical significance is denoted as follows: \**P* < 0.05, \*\**P* < 0.01, \*\*\**P* < 0.001, and **** *P* < 0.0001.

## Results

### The spinal monosynaptic reflex arc becomes segmentally restricted during the first two-weeks of postnatal development

To test for intersegmental supernumerary proprioceptive Ia projections onto a-motor neurons, we used ventral root electrophysiology in the lumbar region of the spinal cord in neonatal mice. We stimulated DRL5 and recorded segmental responses from VRL5 and intersegmental responses from VRL4 (Figure 1A, also see Methods). At P4-7, in addition to a strong segmental response, we also observed a robust intersegmental response (Figure 1B-D). While the amplitude of segmental response was unchanged during development, the intersegmental response began to decline between P8-10, and was lost by the end of second postnatal week (P11-13). Importantly, the intersegmental response is time-locked with the segmental response (Figure 1B). Because we cannot establish monosynapticity using ventral root electrophysiology, we analyzed the onset latencies of the segmental and intersegmental response for each recording. We found that the difference in latencies ranged from 0-300ms (110 ms ± 130 ms), thus supporting the notion that the intersegmental responses recorded in these experiments are monosynaptic (Figure 1E). Indeed, the shortest delay in latency for the onset of possible disynaptic/polysynaptic responses in relation to monosynaptic inputs in motor neurons due to afferent stimulation is in the range of 2-3.5 ms (Mears & Frank, 1997).

Based on the onset latencies of the segmental and intersegmental responses, we predicted the intersegmental response could be monosynaptic. To provide anatomical evidence for direct intersegmental connectivity between L5 proprioceptors and L4 motor neurons, we performed tracing experiments. Application of fluorescent dextrans has been used to label spinal sensory afferents, including proprioceptors (Sürmeli *et al*., 2011; Vukojicic *et al*., 2019), and motor neurons (Vrieseling & Arber, 2006; Kasumacic *et al*., 2010). We visualized DRL5 proprioceptive afferent inputs (yellow, Figure 2) onto VRL4 motor neurons (magenta). We found nearly all L4 motor neurons imaged receive DRL5 proprioceptive inputs at P4-7 (95% ± 4.4%, n = 72 motor neurons, Figure 2A,C). Though the average amplitude of the intersegmental response was 31% of the segmental response, these tracing experiments only reveal the prevalence, not strength, of intersegmental inputs. Thus, based on our functional data, we predict that intersegmental synapses are weaker than segmental synapses, which would be consistent with the observation that they are lost during development (Figure 1B-D). Indeed, by P8-10, we find a significant reduction in the percentage of motor neurons that receive these direct proprioceptive inputs (22.3% ± 6.8%, n = 32 motor neurons, P = 0.0003, Figure 2B,C). This is consistent with a reduction in the intersegmental response amplitude at this age (Figure 1B-D). These anatomical experiments, together with our functional analysis showing intersegmental onset latencies fall within the monosynaptic range, strongly suggests an intersegmental spread of monosynaptic proprioceptive Ia-a-motor neuron synapses. Nevertheless, we will refer to these responses as intersegmental responses in the absence of direct evedince of monosynapticity. Collectively, this shows that while the segmental monosynaptic reflex response persists across early postnatal development, the intersegmental response shows a developmental decline, becoming negligible by the end of the second postnatal week.

**Figure 2.**
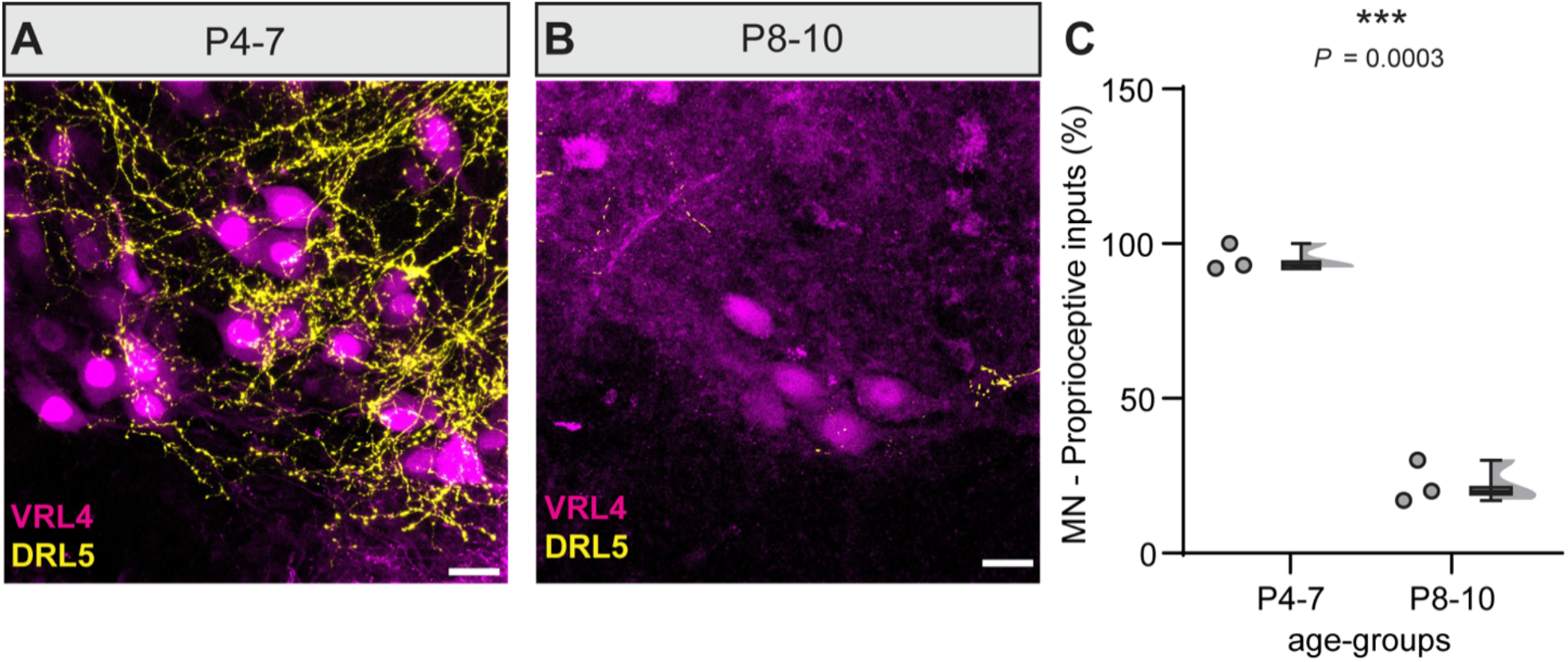
Anatomical validation of direct inputs from L5 proprioceptive inputs onto L4 a-motor neurons. **A** and **B.** Representative 60x confocal (N.A. 1.4) images containing retrogradely labeled L4 motor neurons (Fluorescein, magenta) and proprioceptive inputs from DRL5 (Texas red, yellow) for P4-7 **(A)** and P8-10 **(B).** Scale bars = 20 mm. **C.** Plot showing the percentage of motor neurons that receive proprioceptive inputs. Each dot represents the average prevalence of L5 proprioceptive inputs on L4 motor neurons per section. An average of 24 motor neurons (P4-7) and 11 motor neurons (P8-10) per section were analyzed. Data were analyzes using a Welch’s t test. **A-C** n = 3 sections from N = 3 animals per age-group.

### The monosynaptic reflex arc segmental specification is activity dependent

It is well accepted that synaptic refinement is brought about by activity-dependent mechanisms (Colman *et al*., 1997; Nagappan-Chettiar *et al*., 2024). To determine if loss of the intersegmental response is due to activity-dependent synaptic refinement, we conducted electrophysiological recordings from the Pirt^Cre^;Na_V_1.6^fl/fl^ (Na_V_1.6^cKO^) mouse line, in which we previously found loss of the segmental monosynaptic reflex response by P14 (Espino *et al*., 2025). In our prior work, we analyzed monosynaptic responses from combined ages P6-P11. To determine the precise time course of impairment onset, we conducted new recordings for age groups P8-10 and P11-13 (Figure 3). We found that for P8-10 age group, the segmental response onset latency was significantly slower in Na_V_1.6^cKO^ mice compared to littermate floxed controls (Figure 3A). Additionally, we observed a significant increase in the stimulus threshold required to evoke a segmental response compared to controls (Figure 3B), whereas the monosynaptic peak amplitude was not different between genotypes (Figure 3C). At P11-13, we also found the onset latency was significantly slower (Figure 3D) and stimulus threshold significantly higher (Figure 3E) in Na_V_1.6^cKO^mice compared to floxed controls, and similarly to the P8-P10 age group, the monosynaptic peak amplitude was not significantly different (Figure 3F), though we did note 3 hemicords with negligible amplitudes (0.12 ± 0.06 mV). As in our prior study, by P14-P18 all analyzed parameters were significantly impaired in Na_V_1.6^cKO^ mice compared to controls (Figure 3G-I). Taken together, these data show that the monosynaptic reflex response in Na_V_1.6^cKO^ mice begins to degrade at P8.

**Figure 3.**
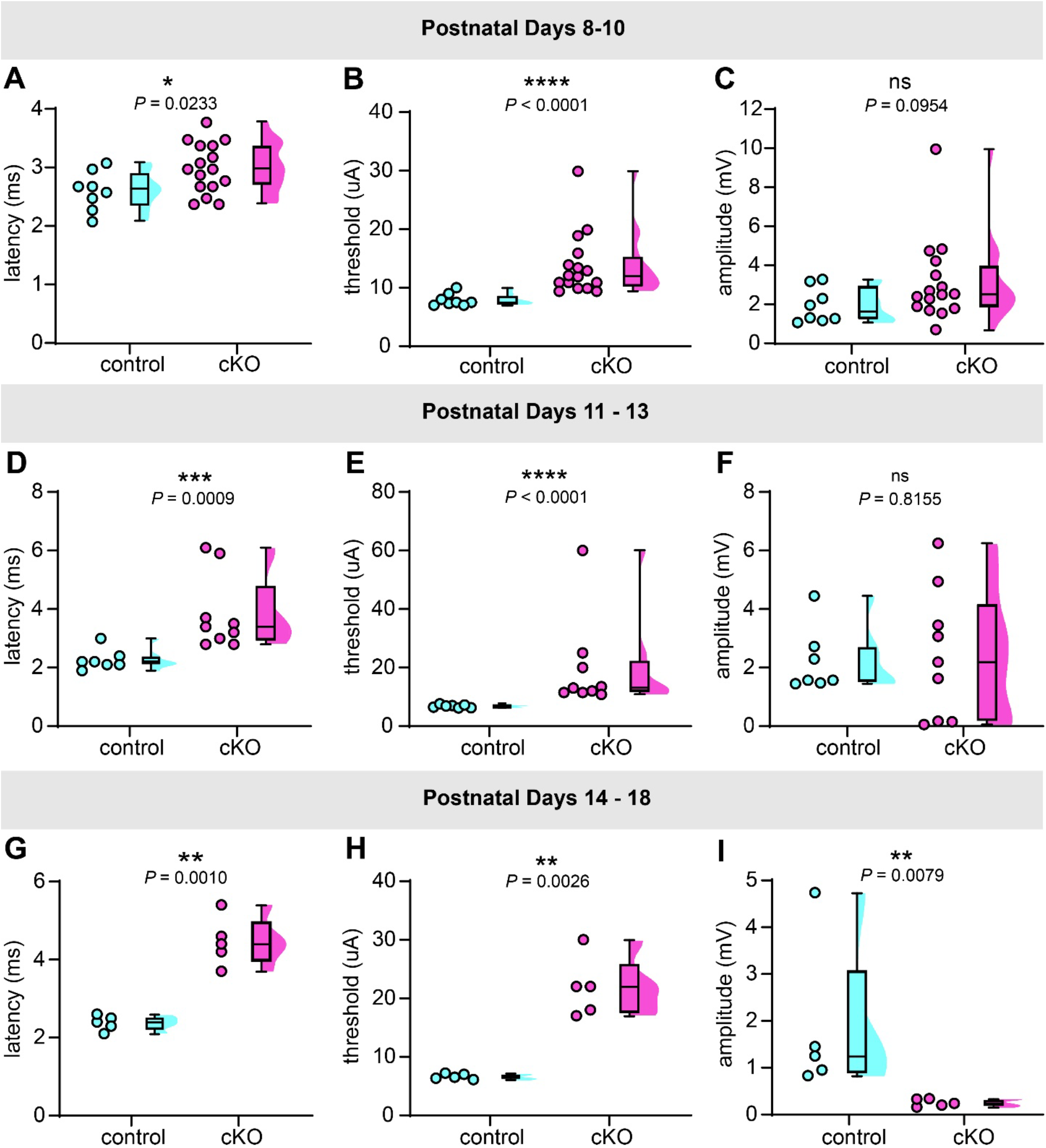
The segmental monosynaptic reflex undergoes progressive developmental degradation in Na_V_1.6^cKO^ mice. **A-C.** Plots showing the onset latency, stimulus threshold and the monosynaptic reflex peak amplitude for Na_V_1.6^fl/fl^ (control) and Na_V_1.6^cKO^ (cKO) mice at P8-10. Statistical analyses were conducted using a Welch’s t-test (**A**) and Mann-Whitney test (**B, C**). Controls: n = 8 hemicords, N = 5 animals, cKO: n = 16 hemicords, N = 10 animals. **D-F.** Plots showing the onset latency, threshold and the monosynaptic reflex peak amplitude for Na_V_1.6^fl/fl^ (control) and Na_V_1.6^cKO^ (cKO) mice at P11-13. A Mann-Whitney test was used for statistical analysis. Controls: n = 7 hemicords, N = 4 animals, cKO: n = 9 hemicords, N = 5 animals. **G-I.** Plots showing the onset latency, threshold, and the monosynaptic reflex peak amplitude at P14-18. Statistical tests included Welch’s t-test (**G, H**) and Mann-Whitney test (**I**). Controls: n = 5 hemicords, N = 5 animals, cKO: n = 5 hemicords, N = 4 animals.

After establishing that the segmental monosynaptic reflex response begins to degrade at P8 in Na_V_1.6^cKO^ mice, we set out to determine if the establishment of segmental specificity of the monosynaptic reflex is altered in an activity-dependent manner. We tested two age-groups, P8-10 and P11-13, and found that the amplitude of the segmental monosynaptic peak for both genotypes was not significantly different (Figures 4A-C, P<0.9999). In line with the findings in C57Bl/6J mice, in control Na_V_1.6^fl/fl^ mice the normalized amplitude of the intersegmental response compared to the segmental response recorded at P8-10 was small (L4/L5: 7.5% ± 4.5 %) and became negligible by P11-13 (L4/L5: 0.3% ± 0.9%). In contrast, Na_V_1.6^cKO^ mice at both P8-10 and P11-13 displayed prominent segmental and intersegmental responses (L4/L5: 25.4% ± 15.1%, 28.6% ± 17.2%, respectively, *P* = 0.6238, Figure 4A-C). As with C57Bl/J6 mice (Figure 1), intersegmental response latencies are time-locked with the segmental response latencies, supporting the notion that the intersegmental responses recorded in Na_V_1.6^fl/fl^ and Na_V_1.6^cKO^ mice could be monosynaptic (Figure 4D). To examine the persistence of intersegmental responses in Na_V_1.6^cKO^ mice anatomically, we again performed tracing experiments as in Figure 2. Like what we observed in P9 C57Bl/6J mice, L4 motor neurons in Na_V_1.6^f/fl^ mice received few projections from L5 proprioceptive Ia afferents (22.8% ± 26.0%, n = 223 motor neurons; Figure 5). Conversely, L4 motor neurons from Na_V_1.6^cKO^ mice received significantly more L5 Ia afferent innervation (64.0% ± 25.4%, *P* = 0.0047; Figure 5B,C), consistent with our electrophysiological recordings. We were unable to see clear axonal labeling near motor neurons as experiments in P4-7 C57Bl/6J mice, likely due to the fragility of axons in Na_V_1.6^cKO^ mice, though clear axonal labeling is visible more dorsally (Supplemental Figure 1). Nevertheless, these results support the notion that the segmental specificity of the monosynaptic reflex arc is activity dependent.

**Figure 4.**
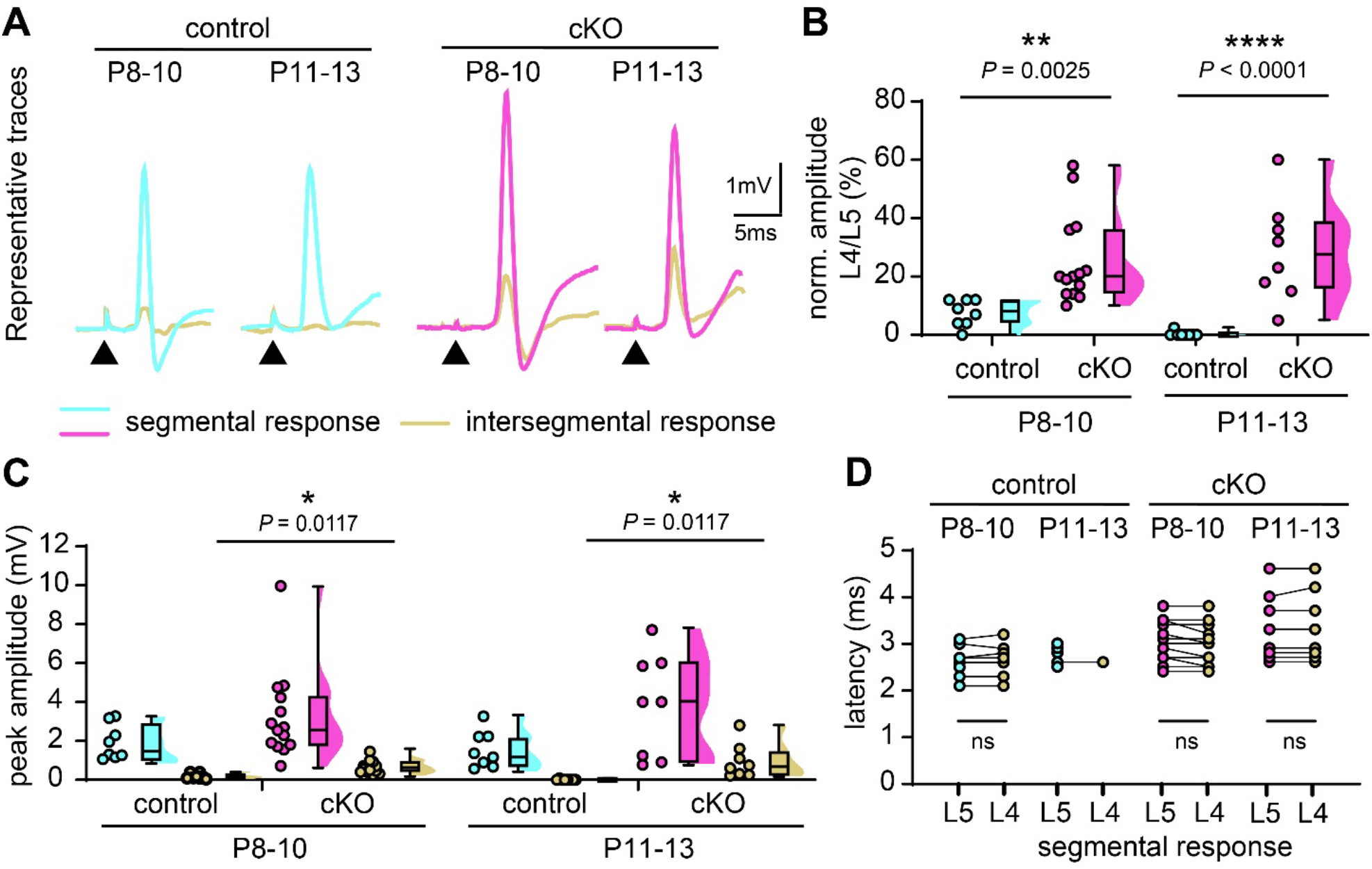
Segmental specificity of the monosynaptic stretch reflex is abolished in Na_V_1.6^cKO^ mice. **A.** Representative traces for the monosynaptic reflex shown. Each pair represents the segmental (L5) and intersegmental response (L4) from the same preparation. Arrowheads indicate onset of stimulation. **B**. Plot showing the intersegmental response peak amplitude (L4) normalized to the segmental response peak amplitude (L5) from the same preparation. Statistical tests included mixed-model two-way ANOVA followed by Uncorrected Fisher’s Least Significant Difference test. **C**. Plot showing the peak amplitude of the monosynaptic stretch reflex. Statistical tests included two-way ANOVA followed by Tukey’s multiple comparisons test. **D**. Plot showing the onset latency of the segmental (L5) and intersegmental response (L4). Straight lines connect pairs of segmental (L5) and intersegmental (L4) values from the same preparation. Statistical tests included Welch’s t-test for comparison between latencies of onset for L5 vs L4 response: P8-10: Controls: *P* = 0.8630, cKO: *P* = 0.6867, P11-13: controls: intersegmental n too small, cKO: *P* = 0.9465. **B - D.** P8-10: Controls: n = 8 hemicords, N = 5 animals, cKO: n = 14 hemicords, N = 10 animals, P11-13: Controls: n = 8 hemicords, N = 4 animals, cKO: n = 8 hemicords, N = 4 animals.

**Figure 5.**
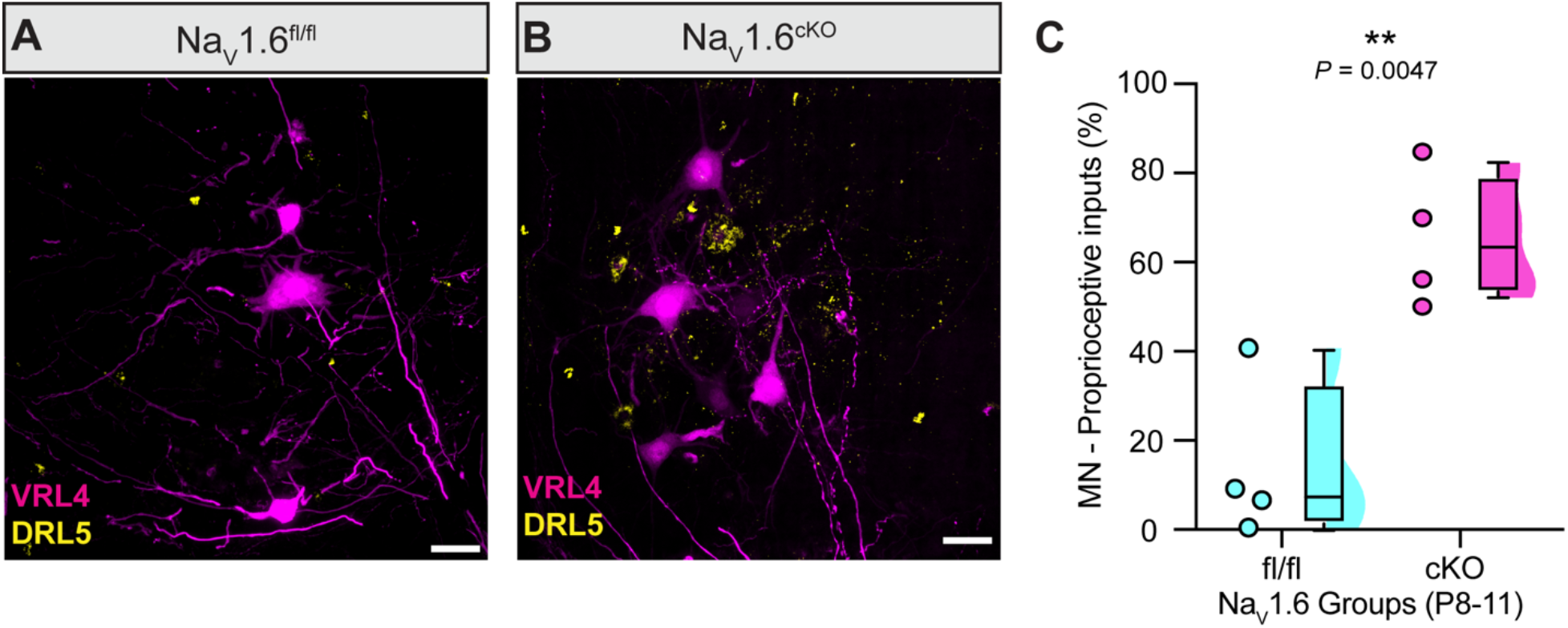
Direct inputs from L5 proprioceptive inputs onto L4 a-motor neurons persist in Na_V_1.6^cKO^ mice. **A** and **B.** Representative 60x confocal (N.A. 1.4) images containing retrogradely labeled L4 motor neurons (Fluorescein, magenta) and proprioceptive inputs from DRL5 (Texas red, yellow) for Na_V_1.6^fl/fl^ **(A)** and Na_V_1.6^cKO^**(B). C.** Plot showing the percentage of motor neurons that receive proprioceptive inputs. Each dot represents the average prevalence of L5 proprioceptive inputs on L4 motor neurons per mouse. A total of 223 motor neurons and 202 motor neurons were analyzed across 4-5 sections from Na_V_1.6^fl/fl^ and Na_V_1.6^cKO^ mice, respectively. Data were analyzes using a Welch’s t test. N = 4 animals per age-group. Scale bars = 25 mm.

### Microglia numbers in the ventral spinal cord are unchanged with impaired proprioceptive activity

To determine whether microglia play a role in the segmental restriction of the monosynaptic reflex arc, we examined their numbers in the ventral spinal cord during the first two weeks of development (Figure 6A-C). We stained spinal cords from C57Bl/6J neonates for the microglia marker IBA1, and ChAT to identify motor neurons, at three different ages: P5, P9 and P12 (Figure 6D-F). We found the number of microglia in the ventral spinal cord showed a significant increase at P9 as well as P12 compared to P5 (Figure 6). This increase in microglial numbers at P9 is in line with the decline in intersegmental monosynaptic reflex amplitude observed during the P8-10 developmental stage (Figure 1). These results indicate that microglia could play a role in establishing segmental specificity of monosynaptic reflex arc during early development. We therefore analyzed microglia numbers near ChAT-expressing motor neurons in spinal cords from P9 Na_V_1.6^fl/fl^ and Na_V_1.6^cKO^ mice (Figure 6G,H); however, there was no genotype dependent difference in microglia counts (Figure 6J), suggesting microglia recruitment is normal in this model.

**Figure 6.**
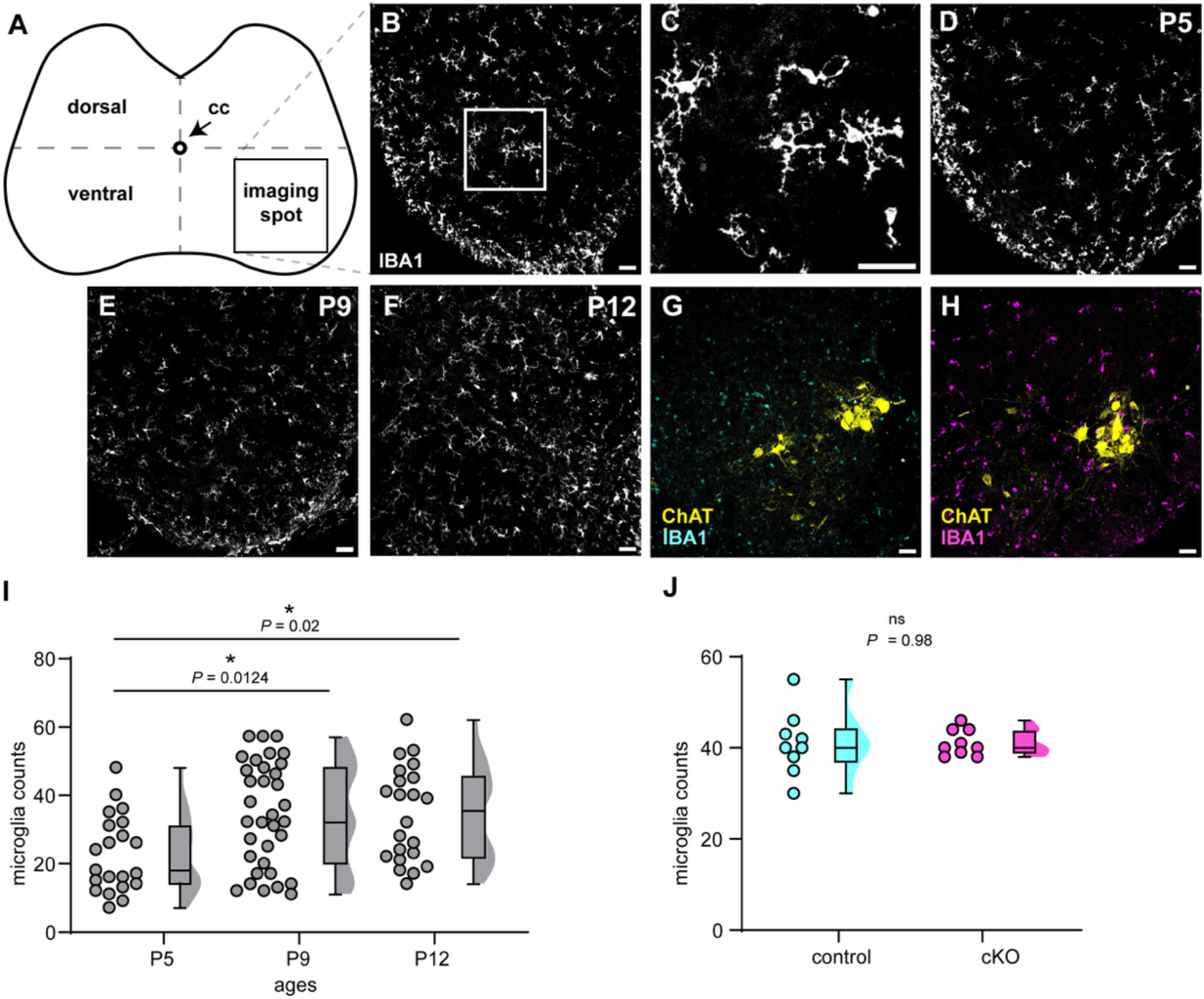
Analysis of microglia numbers in the postnatal ventral spinal cord. Schematic shows a transverse spinal cord section. Lines overlain on the spinal cord passing through the central canal (CC) divide the spinal cord into 4 quadrants. A typical imaging spot in the ventral half of the spinal cord in the bottom right quadrant, is indicated by the square. Image **B** is a representative image containing IBA1+ microglia acquired under a 20X air-objective. Scale bar, 40µm. Image **C** shows an enlargement of the boxed portion from image **B**. **D-F.** Representative images containing IBA1+ microglia acquired under a 20X air-objective for the ages P5 **(D)**, P9 **(E)** and P12 **(F)**, respectively. **G.** Representative image of IBA1+ microglia near ChAT-expressing motor neuron in a spinal cord section from a P9 Na_V_1.6^fl/fl^ mouse acquired under a 20X air-objective. H. Representative image of IBA1+ microglia near ChAT-expressing motor neuron in a spinal cord section from a P9 Na_V_1.6^cKO^ mouse acquired under a 20X air-objective. **I**. Analysis of the number of microglia counts at the indicated ages in the ventral spinal cord of C57Bl/6J mice. Statistical analyses were determined by one-way ANOVA (Kruskal-Wallis test) followed by Dunn’s multiple comparisons test. At P5: n = 21 sections, P9: n = 35 sections, P12: n = 22 sections. N = 3 animals for all ages **J**. Analysis of microglia counts from spinal cords of Na_V_1.6^fl/fl^ (control) and Na_V_1.6^cKO^ (cKO) P9 mice. Controls: n = 9 sections, cKO: n = 9 sections. N = 3 animals for all ages. Statistical significance was determined by a Welch’s t-test. Scale bars, 40 µm

### Impaired proprioceptor activity reduces C1qA expression in the developing spinal cord

Complement signaling is essential for synaptic pruning (Stevens *et al*., 2007; Schafer *et al*., 2012); we therefore investigated if the persistence of an intersegmental response in Na_V_1.6^cKO^ mice could be due to impaired C1q signaling. We carried out immunolabeling for C1qA, the protein that initiates the complement signaling cascade, on transverse sections obtained from the lumbar spinal cord from Na_V_1.6^fl/fl^ and Na_V_1.6^cKO^ mice of both sexes (Figure 7A). Spinal cords were harvested from P9 mice, a time point in which we expected the peak of synaptic refinement to occur based on our initial intersegmental response recordings. Quantification of the mean gray value of the C1qA signal near ChAT-expressing motor neurons showed significantly greater C1qA expression in the ventral spinal cord of Na_V_1.6^fl/fl^ controls compared to Na_V_1.6^cKO^ mice (Figure 7B). This suggests that C1qA signaling is reduced in the ventral spinal cord of Na_V_1.6^cKO^ mice at P9, which is consistent with the persistence of a robust intersegmental response in these mice up to P13.

**Figure 6.**
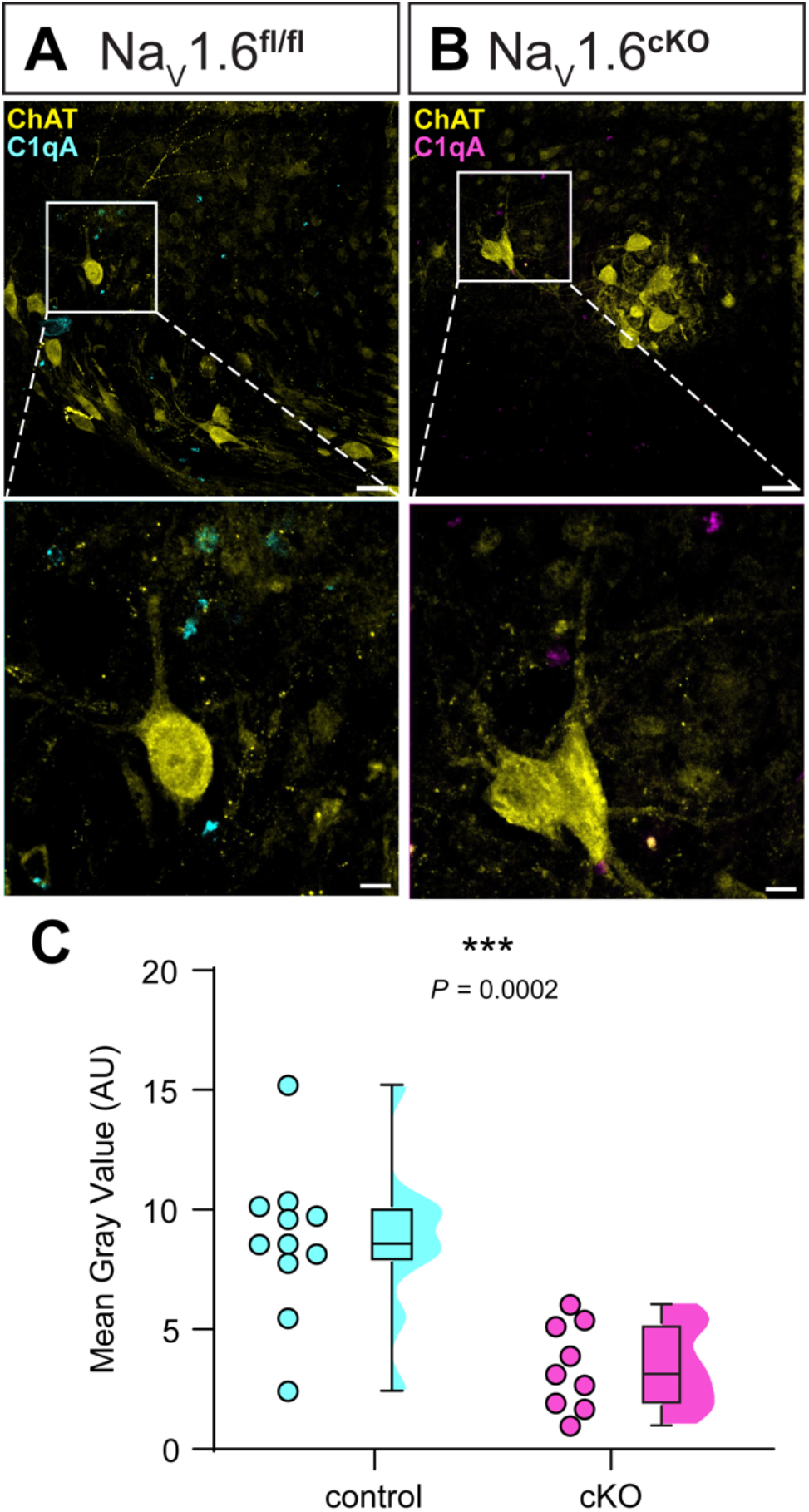
Decreased C1qA expression in the ventral spinal cord of Na_V_1.6^cKO^ mice. **A.** Representative images showing C1qA (magenta) and ChAT-expressing motor neurons (yellow) labeling in transverse sections from ventral spinal cord of control (Na_V_1.6^fl/fl^) and cKO (Na_V_1.6^cKO^) mice. **B.** Plot showing quantification of C1qA expression in the control and cKO mice. Welch’s t-test was used for statistical analysis. Control: n = 11 sections, N = 3 mice; cKO: n = 9 sections, N = 3 mice.

### Segmental monosynaptic reflex is enhanced due to loss of C1qA at P4-7

Reduced C1qA expression in the Na_V_1.6^cKO^ mice prompted us to investigate how loss of C1qA signaling affects the establishment of segmental specificity of the monosynaptic reflex arc. We carried out segmental and intersegmental reflex recordings at P4-7 in C1qA knockout mice (C1qA^- /-^) and littermate controls. We chose this age group to determine if we could recapitulate the presence of a prominent intersegmental response in the C1qA mouse line, as seen in C57Bl/6J mice. Our results show that at P4-7, there is a robust intersegmental response in both controls and C1qA^-/-^ mice (Figures 8A-E). The segmental monosynaptic reflex in C1qA^-/-^ mice had a significantly larger amplitude response compared to the controls (Figures 8B). Moreover, an intersegmental response was present in both genotypes at this age (Figures 8A,B), and the normalized amplitude was not significantly different between the C1qA^-/-^ mice and controls (Figure 7C), despite the segmental monosynaptic reflex amplitude being enhanced in C1qA^-/-^ mice. Notably, the segmental monosynaptic reflex in C1qA^-/-^ mice had a significantly smaller stimulus threshold that gave rise to a larger amplitude response with a significantly shorter onset latency compared to the controls (Figures 8D and E). This finding was surprising, as even at this early timepoint, C1qa^-/-^ mice show an enhancement of every monosynaptic reflex parameter analyzed, possibly indicating a greater exuberance of proprioceptive inputs onto motor neurons compared to aged-matched control mice.

**Figure 8.**
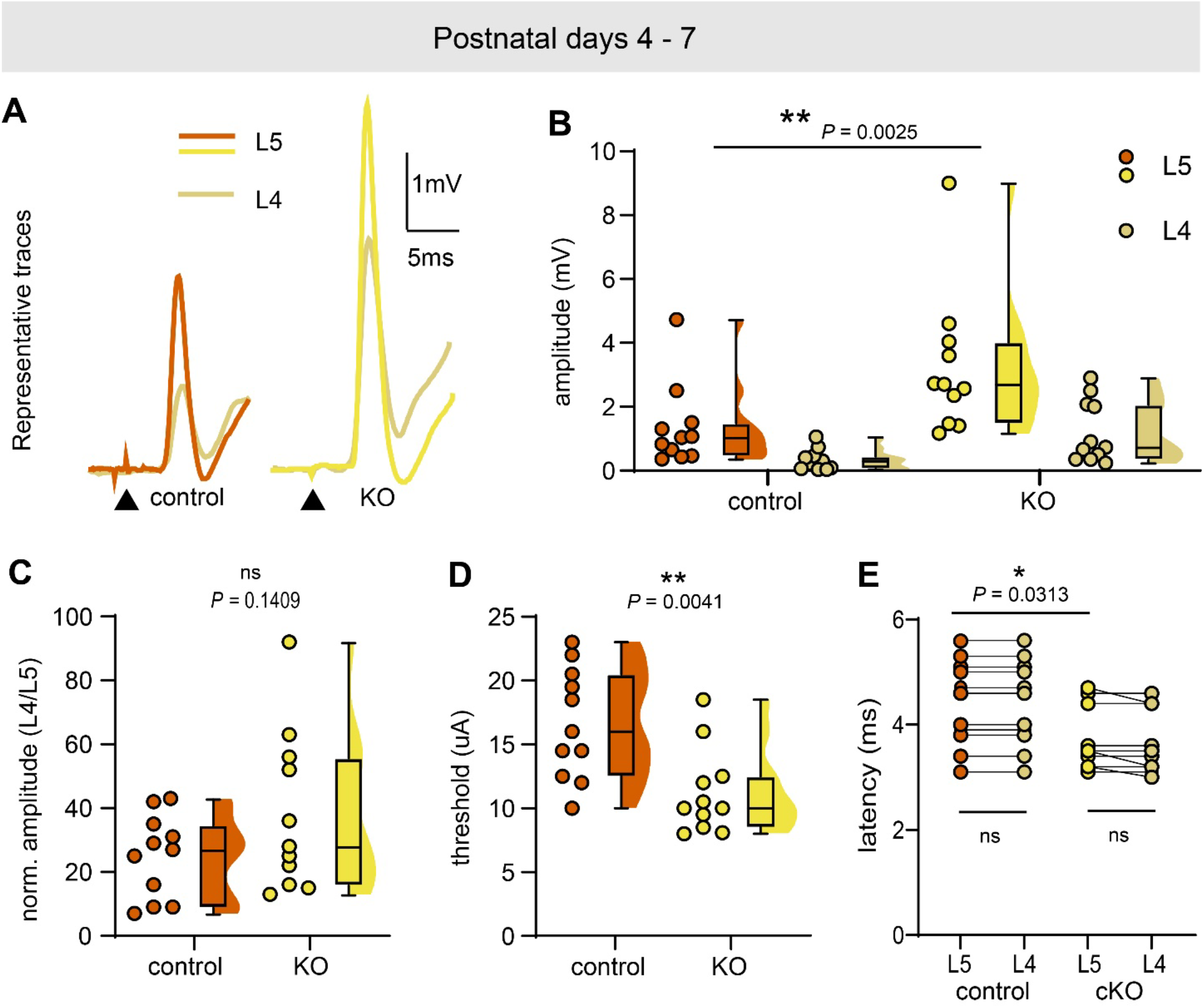
Enhancement of the monosynaptic reflex response in C1qA knockout mice during the first postnatal week. **A.** Representative traces of segmental and intersegmental responses are shown. Each pair represents the segmental (L5) and intersegmental reflex response (L4) from the same preparation. Arrowheads indicate onset of stimulation. L5 and L4 indicate lumbar segments 5 and 4. **B.** Plots showing the peak amplitudes of the segmental monosynaptic and intersegmental reflex response. Statistical tests included a mixed model two-way followed by Uncorrected Fisher’s Least Significant Difference test. **C.** Plot showing the peak amplitude of intersegmental response (L4) normalized to the segmental response (L5) from the same preparation. Statistical tests included Welch’s t-test. **D.** Plots showing the stimulus threshold of the monosynaptic stretch reflex. Statistical tests included Welch’s t-test. **E.** Plots showing the onset latency of the segmental (L5) and intersegmental (L4) response. Statistical tests included a mixed model two-way ANOVA followed by Uncorrected Fisher’s Least Significant Difference test. **B - E.** Controls: n = 11 hemicords, N = 6 animals, cKO: n = 11 hemicords, N = 7 animals.

### Loss of C1qA results in the persistence of an intersegmental response

We next carried out segmental and intersegmental reflex recordings at P11-13 in C1qA^-/-^ mice and littermate controls to determine if the intersegmental response persists at this age range. In line with our hypothesis, at P11-13, C1qA^-/-^ mice show a robust intersegmental response, whereas recordings from littermate control mice showed a negligible intersegmental response (Figures 9A - I), like age-matched C57Bl/6J and Na_V_1.6^fl/fl^ mice. As with the P4-7 age group, the segmental monosynaptic reflex in P11-P13 C1qA^-/-^ mice produced a significantly larger amplitude response with a significantly shorter onset latency that required a significantly lower stimulus threshold compared to controls (Figures 9B, D and E). The normalized intersegmental response amplitude was significantly different between the P11-13 C1qA^-/-^ mice and controls (Figure 9C). While it was negligibly small in the controls (1.4% ± 3.1% of the segmental response), it was 27.1% ± 11.7% of the segmental response in C1qA^-/-^ mice. Thus, genetic deletion C1qA results in loss of segmental specificity of monosynaptic reflex arc in a manner like that of impaired proprioceptor electrical activity.

**Figure 9.**
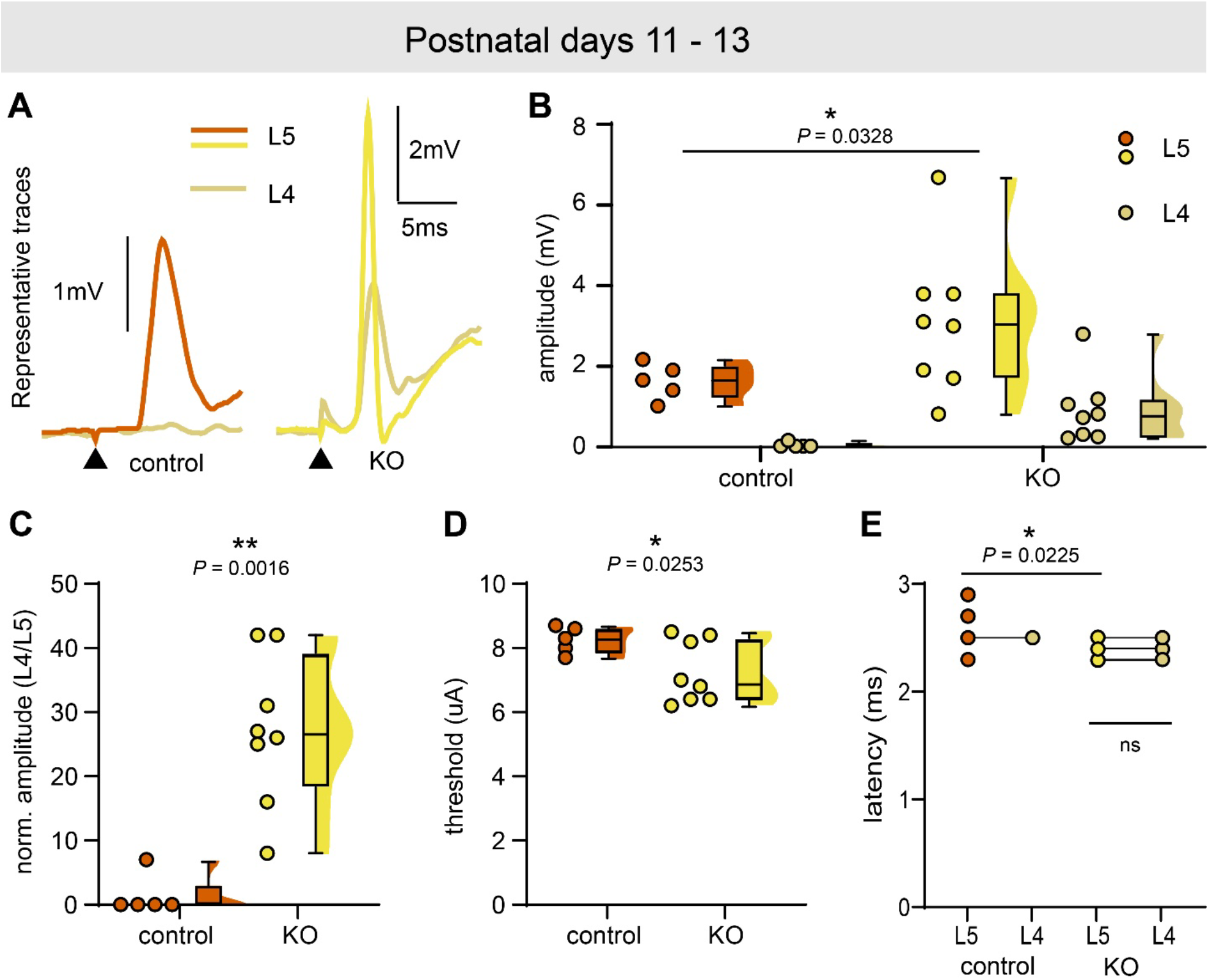
Loss of C1qA results in the persistence of an intersegmental response. **A.** Representative traces of segmental and intersegmental responses are shown. Each pair represents the segmental (L5) and intersegmental reflex response (L4) from the same preparation. Arrowheads indicate onset of stimulation. L5 and L4 indicate lumbar segments 5 and 4. **B.** Plots showing the monosynaptic reflex peak amplitude. Statistical tests included a mixed model two-way ANOVA followed by Uncorrected Fisher’s Least Significant Difference test. **C.** Plot showing the intersegmental response peak amplitude (L4) normalized to the segmental response (L5) from the same preparation. Statistical tests included a Mann-Whitney test. **D.** Plots showing the stimulus threshold of the segmental monosynaptic reflex. Statistical tests included a Welch’s t-test. **E.** Plots showing the onset latency of the segmental (L5) and intersegmental (L4) response. Statistical tests included a mixed model two-way ANOVA followed by Uncorrected Fisher’s Least Significant Difference test. **B-E.** Controls: n = 5 hemicords, N = 4 animals, cKO: n = 8 hemicords, N = 4 animals.

## Discussion

The proprioceptive system and monosynaptic reflex arc have been the subject of numerous investigations, beginning with pioneering studies by Sir Charles Sherrington (Sherrington, 1893). Intersegmental relay of proprioceptive information via spinal interneurons is well established in the cat (Jankowska & Lindström, 1972; Edgley & Jankowska, 1987; Harrison & Riddell, 1989; Jankowska, 1992). Our findings extend this literature by identifying a potentially direct intersegmental connection between Ia afferents and α-motor neurons that is present transiently during early postnatal development and eliminated by complement-mediated pruning. We show that during early postnatal development, in addition to the segmental monosynaptic response recorded at ventral root segment 5 (VRL5) by stimulation of dorsal root segment 5 (DRL5), we also obtained an intersegmental response from the adjacent ventral root (VRL4). This was observed across three different mouse models (C57Bl/6J, Na_V_1.6^fl/fl^, and C1qa^+/+^ mice). In addition to functional studies, we provide anatomical evidence that shows the presence of direct targeting of L4 motor neurons by L5 proprioceptive afferents prior to postnatal day 11. By the end of the second week of postnatal development (P11-13), the intersegmental response was eliminated in C57Bl/6J, Na_V_1.6^fl/fl^, and C1qa^+/+^ mice. Conversely, in Na_V_1.6^cKO^ and C1qA^-/-^ mice, the intersegmental response persisted at least up to P13, the latest developmental timepoint included in our study. We propose a model whereby proprioceptor signaling in the spinal cord leads to activation of the classical complement pathway to prune intersegmental proprioceptive afferents. If proprioceptor function or complement signaling are impaired, supernumerary intersegmental synapses are not effectively pruned (Figure 10). It will be informative to test different mouse models with varying levels of impaired proprioceptor function (Woo *et al*., 2015; Lin *et al*., 2016; Espino *et al*., 2022) to determine if a correlation between the severity of proprioceptor dysfunction and the persistence of an intersegmental response exists.

**Figure 10.**
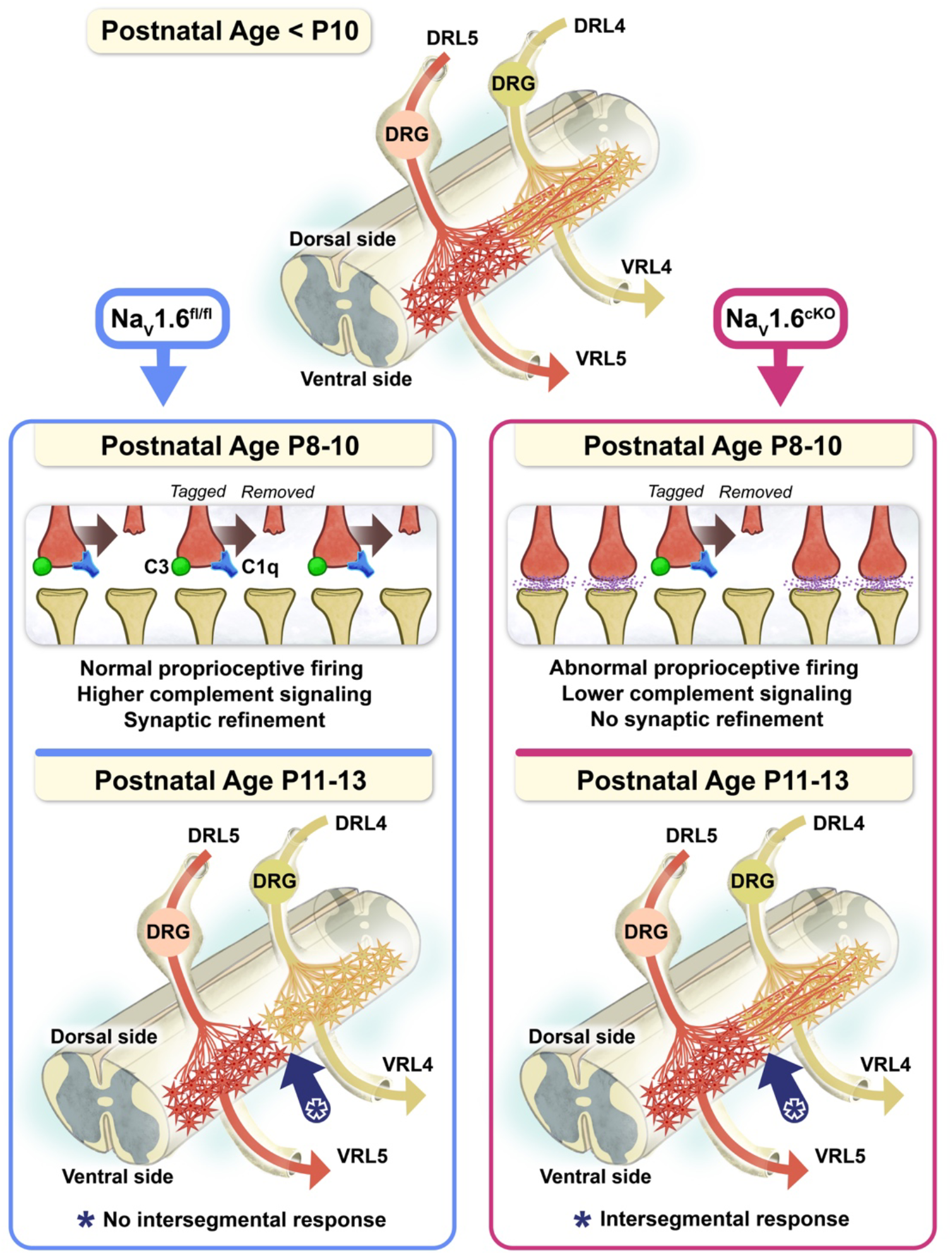
Model of segmental specification of the spinal monosynaptic reflex arc. During the first 10 days of postnatal development, proprioceptive afferents of the dorsal root at lumbar segment 5 (DRL5, red-orange) make synaptic connections onto segmental (red-orange stars) and intersegmental (yellow stars) motor neurons. In control conditions (Na_V_1.6^fl/fl^ shown here), during P8-10, supernumerary intersegmental proprioceptive boutons are tagged with C1qA (blue) initiating the complement signaling cascade and C3 expression (green). The tagged synapses are then pruned, such that by P11-13, no intersegmental response can be evoked. Conversely, in Na_V_1.6^cKO^ mice at P8-10, C1qA expression is low. Thus, fewer intersegmental proprioceptive boutons are tagged, and at P11-13, Na_V_1.6^cKO^ mice retain intersegmental responses.

It is well known that during development, the nervous system undergoes synaptic refinement and elimination of supernumerary synapses (Colman *et al*., 1997; Stevens *et al*., 2007; Schafer *et al*., 2012; Hashimoto & Kano, 2013), which is driven by activity-dependent mechanisms (Huberman *et al*., 2003; Yasuda *et al*., 2021; Faust *et al*., 2021; Nagappan-Chettiar *et al*., 2024). In our experiments using Na_V_1.6^cKO^ mice, we observed the persistence of an intersegmental response in spinal cords of P11-13 animals, which we attribute to reduced synaptic refinement and pruning (Figure 4). During development, inactive or weak synapses are eliminated due to competition with active synapses; conversely, synaptic elimination is impeded if the competition is between two or more weak synapses (Colman *et al*., 1997; Yasuda *et al*., 2021; Faust *et al*., 2021; Nagappan-Chettiar *et al*., 2024). It is conceivable that in Na_V_1.6^cKO^ mice, the segmental monosynaptic reflex could also comprise supernumerary synapses early in development. The concomitant impairments in electrical transmission due to loss of Na_V_1.6, however, makes identifying potential supernumerary segmental synapses difficult. Indeed, deciphering between normal and exuberant synapses is a limitation seen in studies of other neural circuits (Colman *et al*., 1997; Stevens *et al*., 2007; Schafer *et al*., 2012; Hashimoto & Kano, 2013). Nevertheless, based on experiments in C1qA^-/-^ mice, which show an enhanced segmental monosynaptic response, we predict excessive segmental synapses are present, albeit hard to detect, in Na_V_1.6^cKO^ mice.

It is worth noting that some degree of persistent intersegmental connectivity would be predicted even after complete pruning of aberrant connections, because motor pools themselves are not strictly confined to single spinal segments. For example, at the lumbar levels examined here, the tibialis anterior and gastrocnemius motor pools are both found at L3–L4, while the lateral gastrocnemius pool spans L3–L5 (Sürmeli *et al*., 2011). Proprioceptive afferents entering via the L5 dorsal root and forming homonymous connections onto motor neurons of a pool-spanning muscle located in the L4 segment would be expected to persist as ‘correct,’ non-exuberant synapses. Our recordings and tracing, however, were performed at the level of the whole L4/L5 dorsal and ventral roots rather than for individual, molecularly or retrogradely identified motor pools, and we did not observe a corresponding residual physiological or anatomical signal past the second postnatal week. This raises two non-exclusive possibilities: first, the contribution of pool-spanning homonymous connections to the total L5-L4 signal may be small such that its loss falls within the variability of our measurements; second, even pool-spanning homonymous connections may be subject to the same activity-dependent, complement-mediated pruning mechanism we describe. The same could occur for heteronymous inputs. Resolving these possibilities, and whether heteronymous inputs show segmental specificity, would require motor pool-specific labeling (e.g., retrograde tracing from individual muscles) in future work.

The persistence of an intersegmental response in Na_V_1.6^cKO^ mice at P11-13 demonstrates that these synapses, whether homonymous or heteronymous, avoid pruning during postnatal development. We show that in Na_V_1.6^cKO^ mice compared to floxed controls, the segmental monosynaptic reflex response was significantly impaired starting at P8 and continuing until P13, before becoming silent around P14-18, consistent with our prior study (Espino *et al*., 2025). We predict this lack of pruning is due to defective complement pathway activation and reduced synaptic tagging by complement proteins, which in turn impedes the ability of microglia to identify and engulf intersegmental synapses.

Microglia are resident central nervous system immune cells that play an active surveillance role in the nervous system (Nimmerjahn *et al*., 2005; Hanisch & Kettenmann, 2007; Paolicelli *et al*., 2011). Additionally, they play an important role in pruning supernumerary synapses during normal development (Stevens *et al*., 2007; Paolicelli *et al*., 2011; Schafer *et al*., 2012), and their ability to engulf and prune synapses is modulated by sensory experience in the visual system (Tremblay *et al*., 2010). Previous work has also implicated both the classical complement signaling pathway and microglia in excessive pruning of proprioceptive synapses onto a-motor neurons in a mouse model of spinal muscular atrophy (Vukojicic *et al*., 2019). The complement cascade, which is part of the innate immune system, is activated in diseases and disorders as well as during the course of normal development (Stevens *et al*., 2007; Schafer *et al*., 2012). In the current study, we found that in C57Bl/6J mice, microglia numbers are significantly greater in the ventral spinal cord at P9 and P12, compared to P5 (Figure 6), consistent with loss of the intersegmental response during this timeframe. We found no significant difference, however, in microglial numbers between Na_V_1.6^fl/fl^ and Na_V_1.6^cKO^ mice at P9 (Figure 6H). Thus, differences in the number of microglia alone do not account for the persistence of an intersegmental response this mouse line.

Interestingly, we found that expression of a component of the classical complement signaling pathway, C1qA, was significantly reduced at P9 in Na_V_1.6^cKO^ mice compared to control Na_V_1.6^fl/fl^ mice. C1qA tags synapses and induces expression of downstream complement components, such as C3, leading to microglia engulfment of tagged synapses (Dejanovic *et al*., 2018; Wang *et al*., 2020; Chung *et al*., 2025). Furthermore, abolition of complement signaling has been shown to disrupt synaptic refinement during development (Stevens *et al*., 2007; Schafer *et al*., 2012). Thus, we propose that reduced C1qA tagging leads to impaired microglia engulfment of supernumerary intersegmental synapses, resulting in their persistence and an evocable intersegmental response in Na_V_1.6^cKO^ mice (Figures 4 and 10). This is consistent with the persistence of an intersegmental response in C1qa^-/-^ mice through the second week of postnatal development (Figures 9). These data also suggest that normal electrical activity in proprioceptors is essential for appropriate levels of C1qA expression in the ventral spinal cord. It has previously been shown that retrograde signaling from proprioceptive afferent terminals is required to differentiate presynaptic inhibitory terminals in the spinal cord for effective presynaptic inhibition (Mende *et al*., 2016). It is possible that similar paracrine signaling mechanisms trigger C1qA release from microglia for synaptic tagging and subsequent refinement. Nevertheless, the mechanisms underlying how proprioceptive signaling induces C1qA expression in the ventral spinal cord, and the identity of potential paracrine signaling molecules, remain to be identified.

A limitation of our study is the lack of a functional demonstration that shows the intersegmental response is a *bona fide* monosynaptic connection between DRL5 proprioceptive afferents and VRL4 a-motor neurons. We conducted two specific analyses to address this limitation. First, we compared the onset latencies of the segmental and intersegmental responses (Figures 1,4,8 and 9) to show they are time-locked and within monosynaptic latencies (110 ms ± 130 ms, Mears & Frank, 1997). Second, we employed anatomical tracing that showed direct proprioceptive L5 afferent inputs onto traced L4 a-motor neurons (Figure 2 and Figure 5). Based on these experiments, we predict the intersegmental response is monosynaptic. Despite this limitation, our experimental protocol (see Methods) provides a means of selectively targeting and studying intersegmental supernumerary synapses and exploring the mechanistic basis of their synaptic refinement. This work also provides a new model for studying nervous system refinement during development.

Movement is driven by muscle contraction. Each muscle is innervated by a distinct group of α-motor neurons known as a motor pool, which is regulated by upstream premotor interneurons. To uncover the organizational logic of these circuits, studies have utilized muscle-specific injections of CTB (Surmeli et al., 2011) or monosynaptically restricted, trans-synaptic rabies virus tracing to map premotor networks along the spinal cord (Tripodi *et al*., 2011). While a motor pool targets a single muscle, its axons exit the spinal cord through one or more ventral roots. We believe fully deciphering motor functionality requires integrating this segment-specific organization with pool-specific logic. Taken together, our data demonstrate that the proprioceptive monosynaptic reflex arc is segmentally restricted by the end of the second postnatal week through a developmental program driven by proprioceptive signaling that activates the complement cascade to eliminate exuberant intersegmental synapses.

## Additional information section

### Data availability statement

All original ‘raw’ data presented in graphical form for this manuscript has been archived and will be made fully available upon reasonable request.

### Competing Interests

None to declare

## Author Contributions

CN: Conceptualization, data curation, formal analysis, investigation, methodology, project administration, supervision, validation, visualization, writing-original draft, writing - reviewing and editing; SO: Data curation, formal analysis, investigation, visualization, writing - reviewing and editing; ARM: Data curation, formal analysis, investigation, writing - reviewing and editing; AS: Data curation, formal analysis, investigation, writing - reviewing and editing TNG: Conceptualization, funding acquisition, project administration, resources, supervision, visualization, writing - reviewing and editing.

## Artificial Intelligence Generated Content (AIGC) policy

Anthropic Claude Opus 4.8 was used for review and editing. The original draft was written by authors without AI assistance.

## Acknowledgments

We thank Joshua Philip Tulman for the summary figure. We would also like to thank the Griffith and Contreras Lab members for helpful discussions.

## Funding

This study was supported by the National Institute General Medical Sciences (2T32GM135741 and 5T32GM153586, S.O.), National Institute of Neurological Disease and Stroke (R01NS135005, T.N.G), a Sloan Research Fellowship (FG-2024-21522, T.N.G), a McKnight Scholar Award, and a Howard Hughes Medical Institute Freeman Hrabowski Scholar Award.

---

**Supplemental Figure 1.**
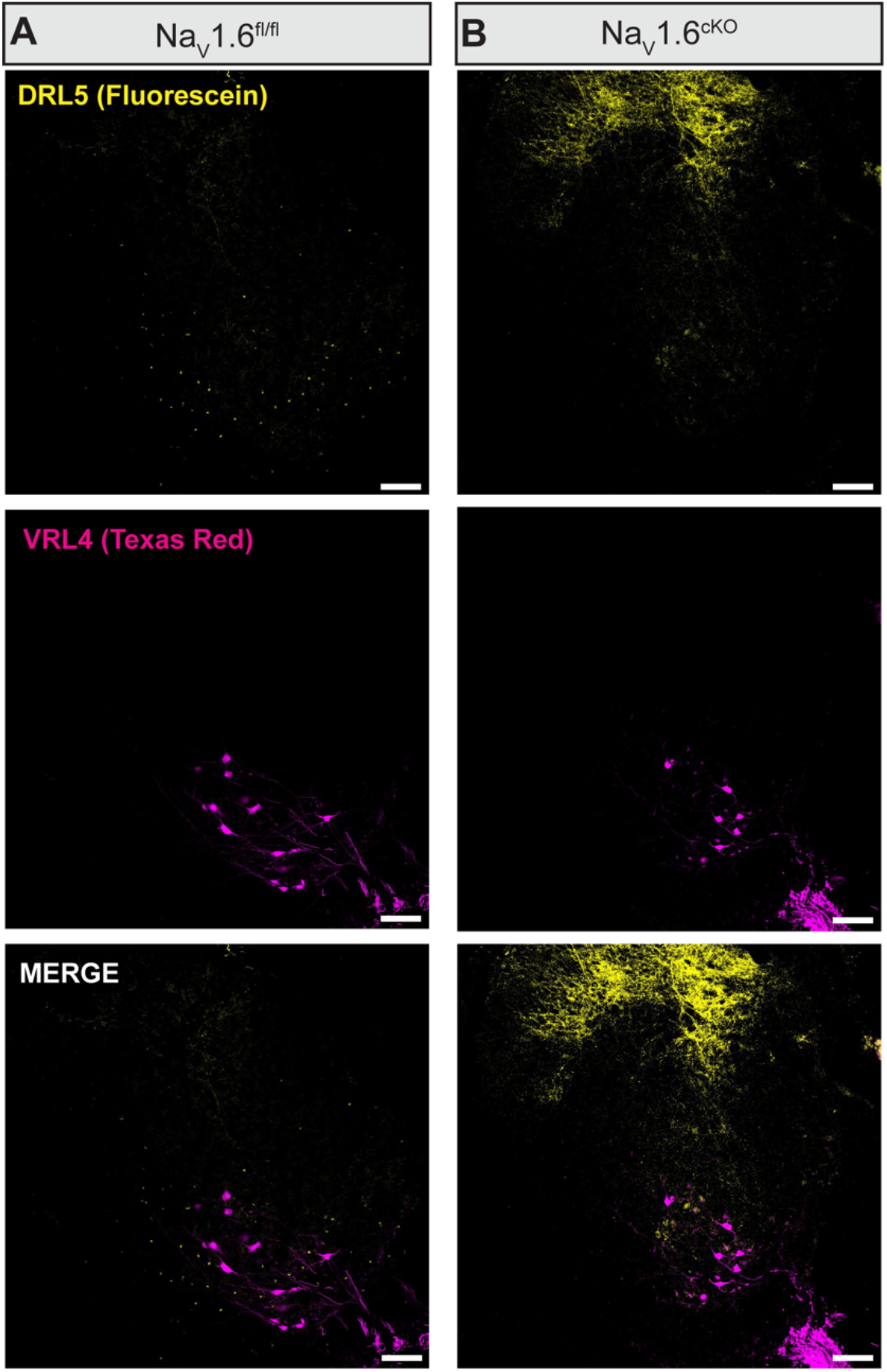
A broad increase in sensory innervation of the L4 spinal segment by L5 doral root axons in Na_V_1.6^cKO^ mice. Representative 10x confocal (air objective) images containing retrogradely labeled L4 motor neurons (Fluorescein, magenta) and sensory axons from DRL5 (Texas red, yellow) from Na_V_1.6^fl/fl^ **(A)** and Na_V_1.6^cKO^**(B)** mice.

## Notes

### Competing Interest Statement

The authors have declared no competing interest.

### Summary of Updates

This version of the manuscript has been revised to include additional citations in the introduction and discussion, as well as new anatomical data showing proprioceptive inputs from dorsal root L5 persist on motor neurons at the spinal segement L4. This data is now in a new Figure 5, as well as a Supplemental Figure 1.

